# Isolation disrupts social interactions and destabilizes brain development in bumblebees

**DOI:** 10.1101/2021.12.16.472962

**Authors:** Z Yan Wang, Grace C. McKenzie-Smith, Weijie Liu, Hyo Jin Cho, Talmo Pereira, Zahra Dhanerawala, Joshua W. Shaevitz, Sarah D. Kocher

**Affiliations:** Department of Ecology and Evolutionary Biology, Princeton University, Princeton, NJ; Lewis Sigler Institute of Integrative Genomics, Princeton University, Princeton, NJ; Department of Physics, Princeton University, Princeton, NJ; Princeton Neuroscience Institute, Princeton University, Princeton, NJ; Department of Neuroscience, Washington University School of Medicine, St. Louis, MO, USA

## Abstract

Social isolation, particularly in early life, leads to deleterious physiological and behavioral outcomes. Few studies, if any, have been able to capture the behavioral and neurogenomic consequences of early life social isolation together in a single social animal system. Here, we leverage new high-throughput tools to comprehensively investigate the impact of isolation in the bumblebee (*Bombus impatiens*) from behavioral, molecular, and neuroanatomical perspectives. We reared newly emerged bumblebees either in complete isolation, small groups, or in their natal colony, and then analyzed their behaviors while alone or paired with another bee. We find that when alone, individuals of each rearing condition show distinct behavioral signatures. When paired with a conspecific, bees reared in small groups or in the natal colony express similar behavioral profiles. Isolated bees, however, showed increased social interactions. To identify the neurobiological correlates of these differences, we quantified brain gene expression and measured the volumes of key brain regions for a subset of individuals from each rearing condition. Overall, we find that isolation increases social interactions and disrupts gene expression and brain development. Limited social experience in small groups is sufficient to preserve typical patterns of brain development and social behavior.

## Results and Discussion

Social animals rely on interactions with conspecifics to survive. Isolation from the social group leads to detrimental impacts on physical health, fitness, and even longevity^1–6^. The effects of social isolation are even more profound during sensitive developmental periods, such as in early life, when social experiences may strongly influence an individual’s “social competence”, the ability to adapt behavior according to changes in social context^7–9^. This can lead to poorer developmental or fitness outcomes^10^. For example, increased aggression across social contexts is a common consequence of social isolation in mice^11,12^, fish^8,13^, flies^14–16^, and crickets^17,18^.

The early life environment may also impact social competence in the social insects, who live collectively in colonies ranging from a few individuals to millions^19^. A growing body of research shows that social isolation impacts the behavior and physiology of bees^20–22^, ants^1,2,4,23,24^, and wasps^25–27^. Few studies have been able to capture behavioral and neurogenomic consequences of early life social isolation in a single social animal system. Here, we investigate the impacts of social isolation in the bumblebee (*Bombus impatiens*) on individual and social behavior, gene expression, and neuroanatomy.

Bumblebees live in social colonies consisting of about 100-200 female workers and a single queen^28^. Within the colony, individuals display consistent differences in behavior that are stable over time and context. These behavioral repertoires are established in the first 1-2 weeks of adulthood through pairwise and spatial interactions among individuals ^29–32^. During this same period in early adulthood, the bumblebee brain is rapidly developing^33^. To determine if and how social isolation impacts social behaviors in this species, we experimentally altered the social environments of workers during this early life developmental period, and we assayed individual bee behavior either alone or paired with a social partner.

To alter the early life social experiences of bumblebees, we developed a modular housing chamber to isolate residents from external auditory, visual, and odor cues (Methods, Figure 1A). We collected newly-eclosed callow females, recognizable by their silvery appearance and sluggish behavior in the colony^28^, and split them amongst 3 different early life treatment conditions: isolation (Iso, n = 96 for behavior, n = 16 for RNA sequencing, n = 20 for imaging), in which a single bee is housed in complete social isolation; group-housed (Grp, n = 113 for behavior, n = 15 for RNA sequencing, n = 24 for imaging), in which four nestmates are co-housed outside the colony; and colony-housed (Col, n = 99 for behavior, n = 9 for RNA sequencing, n = 22 for imaging), in which the individual is immediately returned to her natal colony (Figure 1A). Bees were kept in their treatment condition for 9 consecutive days, thus isolated bees were reared completely devoid of social experiences and group- and colony-housed bees experienced varying amounts of socialization. On post-eclosion day 10, behavioral assays were performed and tissues were collected for downstream analysis. 306 bees were included in the behavioral trial assays, 40 bees in the transcriptomic analyses, and 66 bees in the volumetric analyses (see Methods for details). Different sets of individuals were used for each downstream analysis (behavior, brain gene expression, brain morphology), precluding analyses that combined multiple datasets.

**Figure 1.**
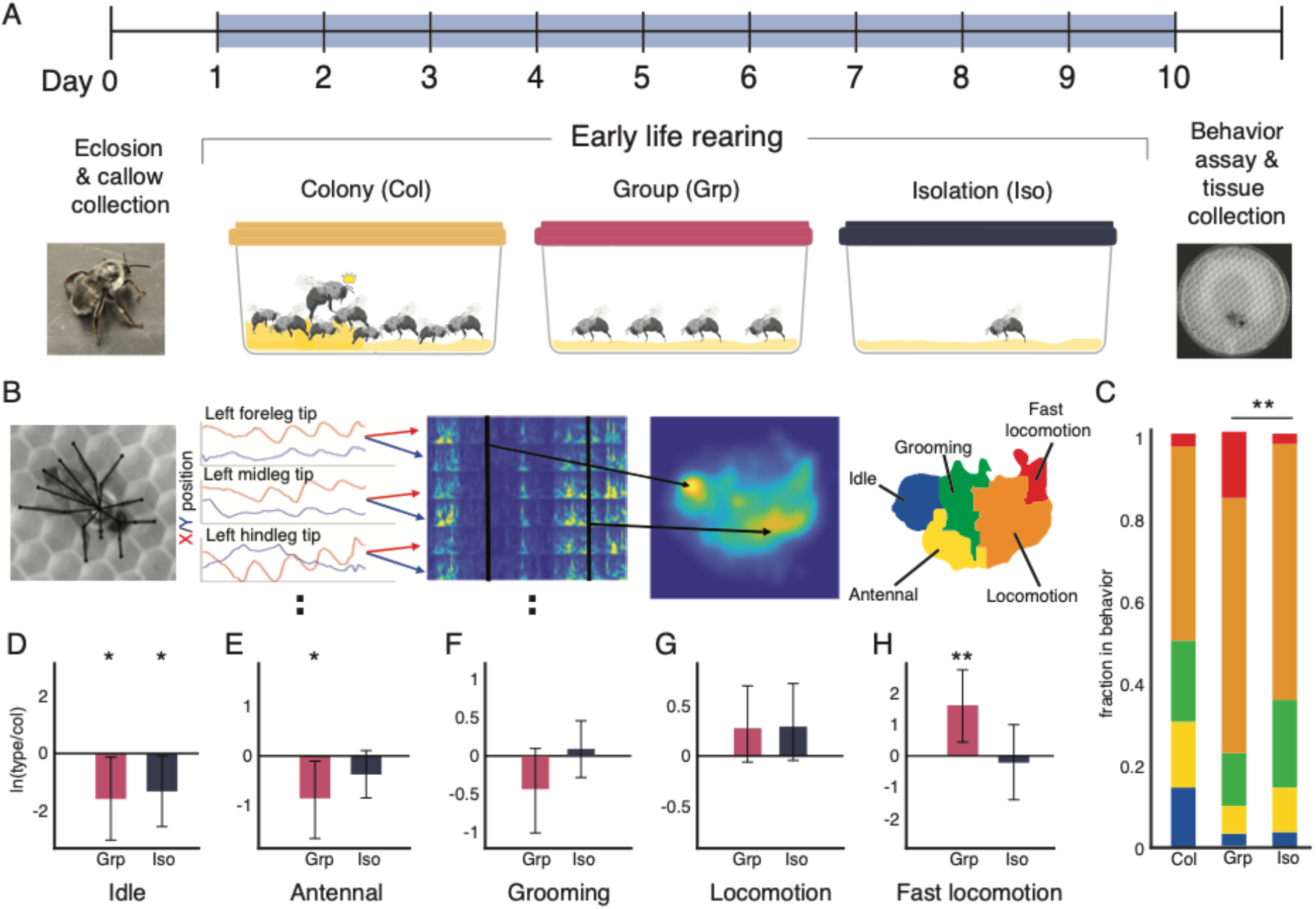
Early life rearing condition alters adult behavior in the bumblebee. **A.** Experimental overview. Newly emerged callows, identified by their silver-white pigmentation and slow, sluggish gait, were assigned to one of three treatment conditions: colony (col) in which the individual is returned to her natal colony; group (grp), in which four nestmates are co-housed outside of the colony; and isolation (iso), in which a single bee is housed in complete social isolation. Bees were housed in these treatment conditions for 9 consecutive days. On post-eclosion day 10, bees were collected for behavioral and neurobiological assays. **B.** Behavioral assay and embedding. Freely-behaving bees were recorded from above for 30 minutes under IR illumination. Bees were assayed in either solo or paired contexts. SLEAP was used to track body parts and bee identity over the duration of the behavioral assay (black overlay). The spectrograms of body part time traces were embedded into a two-dimensional space using t-SNE. Regions of high density were clustered using a watershed transform, then grouped together according to their common behavior motifs into a behavior map of 5 discrete behaviors: idle, antennal behaviors, grooming, locomotion, and fast locomotion. For full details, see Methods. **C**. Time use compositions. Isolated and group-reared bees differ significantly in their time usage compositions (nonparametric multivariate test on ilr transformed fractions, Wilks’ Lambda type statistic. ** indicates p < 0.01). **D-H**. Compositional analysis of discrete behaviors. There are significant differences across solo bees in the idle, antennal, and fast locomotion behaviors as quantified by the log ratio differences between geometric means of iso or grp bees vs. col bees (error bars are bootstrapped 95% confidence intervals). Differences are considered significant in the bootstrapped 95% confidence intervals do not overlap 0. *p<0.05, **p<0.01.

We captured the behavior of experimental bees by themselves (“solo”) and with another bee (“paired”) by recording their free behavior in a 10-cm petri dish for 30 min under infrared illumination, which bees cannot see^34^. For paired conditions, we assayed same-treatment pairs (Iso+Iso, Grp+Grp, Col+Col) as well as each possible combination (Iso+Grp, Iso+Col, Grp+Col, and Grp bees from separate groups) (Figure S1-2). To quantify the behavior of the bees, we used the MotionMapper technique^35^. We first used the SLEAP^36^ pose tracking software to identify body parts in each frame (Figure 1B, Video S1), and then performed a continuous wavelet transform on the body part position time series. The concatenated spectral densities were then embedded into two dimensions using t-distributed stochastic neighbor embedding (t-SNE), and a probability density function of time spent at each location of the t-SNE space revealed peaks corresponding to commonly repeated body part dynamics (Figure 1B, Figure S1, see Methods for details). We segmented the embedded space via a watershed transform to separate regions of stereotyped limb dynamics. We assigned each region to one of five discrete behavior states based on corresponding video clips: idle (no movement), antennal movement, grooming, locomotion, and a fast locomotion behavior mostly seen in solo trials of group-reared bees (Figure 1, Figure S1, Video S2). This enabled us to define discrete behavior composition profiles for each bee and pinpoint behavioral biases in each treatment group. These analyses reveal that colony-reared bees spend more time in an idle state than group-reared or isolated bees, and group-reared bees spend much more time in fast locomotion than colony-reared or isolated bees (Figure 1C). These data are supported by quantification of the bees’ instantaneous speeds over the course of the behavior assay (Figure S1E).

We then used principles from compositional data analysis to quantify differences in the behavioral profiles of each treatment group in the absence of a social partner^37^. We carried out an isometric log-ratio (ilr) transform on the fraction of time spent in each behavioral state to express the data in terms of four independent components. We then performed a non-parametric multivariate analysis on the ilr components. This analysis reveals a significant difference between the overall behavioral profiles of the isolated and group-reared bees when alone (Wilks’ Lambda type statistic, p<0.01, Figure 1C). To examine if and how behaviors differ between the colony-reared (control) and group-reared or isolated bees, we calculated the log-ratio of the geometric means for each behavioral state across individuals. We find that isolated and group-reared bees spend less time in an idle state than colony-reared bees do, and that group-reared bees spend more time in fast locomotion and less time on antennal behaviors (a primary mode of communication in bees)^28^ compared to colony-reared bees (Figure 1D-H; see Methods for details). These data highlight the impact of early life environments on individual behavior: in solo contexts, bees from all three treatment groups diverge in behavior in unique ways.

Next, we examined the differences in behavior among treatment groups in the presence of a social partner because pairwise interactions are considered the building blocks of group behavioral dynamics^38^. We first quantified how often paired bees were in close proximity by examining the differences of inter-thorax distance distributions compared to random chance (see Methods) (Figure 2A). For clarity, we present the results of only the same-treatment pairs, but all pairwise comparisons are presented in the Supplement (Figure S2). We found all pairings to be enriched for inter-thorax distances less than 2 cm. To determine whether distance from a social partner impacts a bee’s overall behavioral repertoire, we quantified changes in the behaviors of paired bees depending on their distance from a social partner using the Jensen-Shannon divergences between 0.2 cm-binned limb dynamics (i.e. the average t-SNE embedded spaces of bees) and the limb dynamics at 8 cm (see Methods)^39^. We find that, across all pairing types, behavior changes strongly when the bees are less than 2 cm (roughly two body-lengths) apart but that the bees’ behavior is largely unaffected by the partner at larger distances (Figure 2B). Based on these results, we defined bees to be affiliated when their inter-thorax distance is less than 2-cm and unaffiliated when they are farther apart. This is concordant with previous studies defining social interactions in a similar range^32^. Surprisingly, we found that pairs of isolated bees spent the most time affiliated with social partners across all pairings.

**Figure 2.**
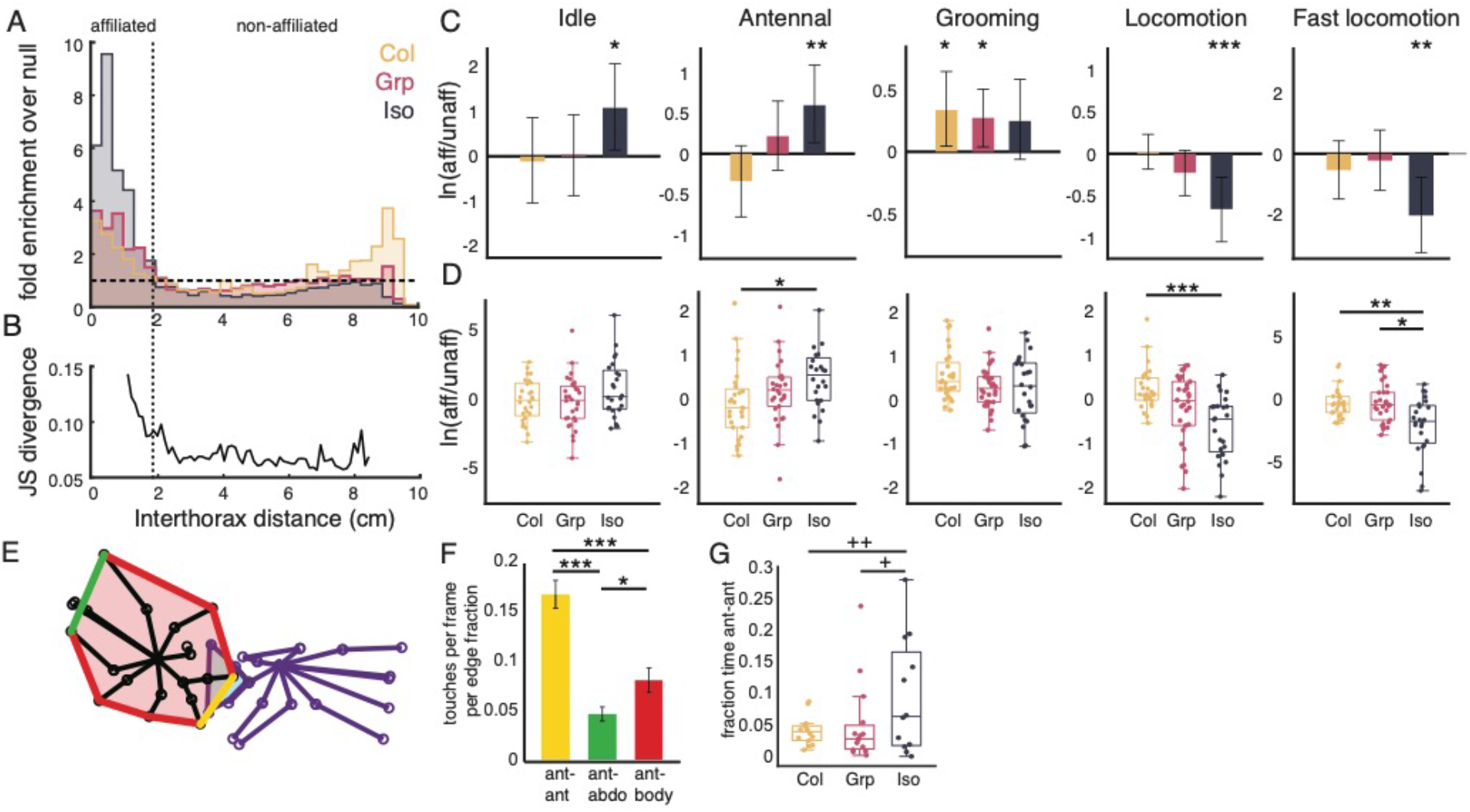
Isolated bumblebees display altered social interactions. **A**. Bees are considered to be affiliated at inter-thorax distances of 2cm or less (within the vertical dotted line). Pairs of isolated, group-, and colony-reared bees are enriched within this distance compared to the null (occupancy above horizontal dashed line, which indicates expected level for randomly arranged bees). **B**. The difference in overall behavior map calculated at each inter-thorax distance and then compared to a ‘far’ distance of 8cm using Jensen-Shannon divergence indicates a difference in behavior that drops off after 2cm. **C**. Isolated bees spend significantly more time in the idle and antennal states, and less time in the locomotion and fast locomotion states when affiliated, as indicated by the log ratio of the geometric means of time spent in each behavior. Differences are considered significant in the bootstrapped 95% confidence intervals do not overlap 0. *p<0.05, **p<0.01, ***p<0.001. **D**. Pairs of isolated bees differ from pairs of colony bees in the change in occupancy of the antennal state when affiliated, and differ from both pairs of colony bees and pairs of grouped bees in the change in occupancy of the locomotion and fast locomotion states when affiliated (Kruskal-Wallis test, Wilcoxon rank sum for pairwise comparisons with Bonferroni correction, *p<0.05, **p<0.01, ***p<0.001). **E**. Antennal (yellow), abdominal (green), and body (red) edges define where antennation occurs. **F**. Pairs of colony bees engage in more antennae-to-antennae touches per available edge space compared to antennae-to-abdomen or antennae-to-body touches (Kruskal-Wallis test, Wilcoxon rank sum for pairwise comparisons with Bonferroni correction, *p<0.05, **p<0.01, ***p<0.001). **G**. Pairs of isolated bees have significantly more variance in the amount of time they spend antennae-to-antennae than group- or colony-reared bees (nonparametric Fligner-Killeen test with Bonferroni correction, +p<0.05, ++p<0.01).

We then compared the behavioral profiles of bees when affiliated versus unaffiliated (Figure 2C). Affiliation has the broadest impact on the behaviors of isolated bees: isolated bees locomote less and engage in more idle and antennal behaviors when they are close to a social partner. In contrast, only grooming is statistically affected by affiliation in the group- and colony-reared bees, where it is increased during affiliation (Figure 2C). Comparing across rearing conditions, we find that affiliation has a statistically different effect on the antennal, locomotive, and fast locomotive behaviors of isolated bees compared to colony-reared bees, and on fast locomotive behavior of isolated bees compared to group-reared bees. There is no such difference between the group- and colony-reared bees (Figure 2D). Taken together, our results reveal that isolated bees not only spend more time close to their social partners, but their behaviors are also more strongly impacted by partner proximity than any other rearing condition.

Finally, we characterized the differences in antennation behaviors across the rearing conditions (Figure 2E-G). Affiliation and other modes of social cooperation often rely on the ability to discriminate between nestmates and non-nestmates. Bumblebees and other social insects primarily do this with chemical signals they detect via chemosensory receptors on their antennae^40,41^. The chemical composition of these signals can vary across body parts^42,43^, so where the antennae make contact may indicate targeting a particular subset of odorants over others. To determine which mode of antennation has the most relevance in this context, in the colony-reared (control) bees, we normalized antennal touches to the antennal, rear abdominal, and body zones of the partner bee to account for differences in the size of each zone (Figure 2E).

We find that pairs of colony-reared bees have significantly more antennae-to-antennae touches than antennae-to-abdomen or antennae-to-body (Figure 2F), demonstrating that antennae-to-antennae touching is a favored mode of contact over what is expected by random chance. We then compared antennae-to-antennae touches across all rearing conditions, and find that all bees spent comparable fractions of time engaging in antennae-to-antennae touching (Figure 2G). However, isolated bees showed significantly higher variance in their propensity for antennae-to-antennae touching compared to colony- and group-reared bees, suggesting that this behavior may be modulated by early life social experience.

Together, the results from our behavioral assays demonstrate that the early life social environment induces changes in key social behavioral features later in life. Both isolated and group-reared bees showed perturbed behavioral profiles in solo assays compared to colony-reared bees. However, in paired assays, isolated bees have broad and significant changes to their behavior when affiliated with a partner bee. In contrast, the behaviors of both group- and colony-reared bees are largely unaffected by proximity to a social partner. Isolated bees also show a large variance in the amount of time they spend in antennae-to-antennae contact with a partner bee, while group- and colony-reared bees are more uniform. This suggests that, while the extra-hive environment of the group-rearing condition alters the behavior of bees when they are alone, only the isolated bees have perturbed behavior in the presence of a social partner.

The behavioral differences we identified between isolated and group-reared bees suggests that there may be underlying neurobiological differences between these experimental groups. To better understand the molecular underpinnings of these behavioral changes, we performed whole brain transcriptome sequencing on a subset of treatment bees (isolated, n=16; group, n=15; colony, n=9) using TM3’seq, a tagmentation-based 3’-enriched RNA sequencing approach^44^. We first performed an analysis of differentially expressed genes (DEGs) across treatment groups, blocking for natal colony (Figure 3A, see Methods).

**Figure 3.**
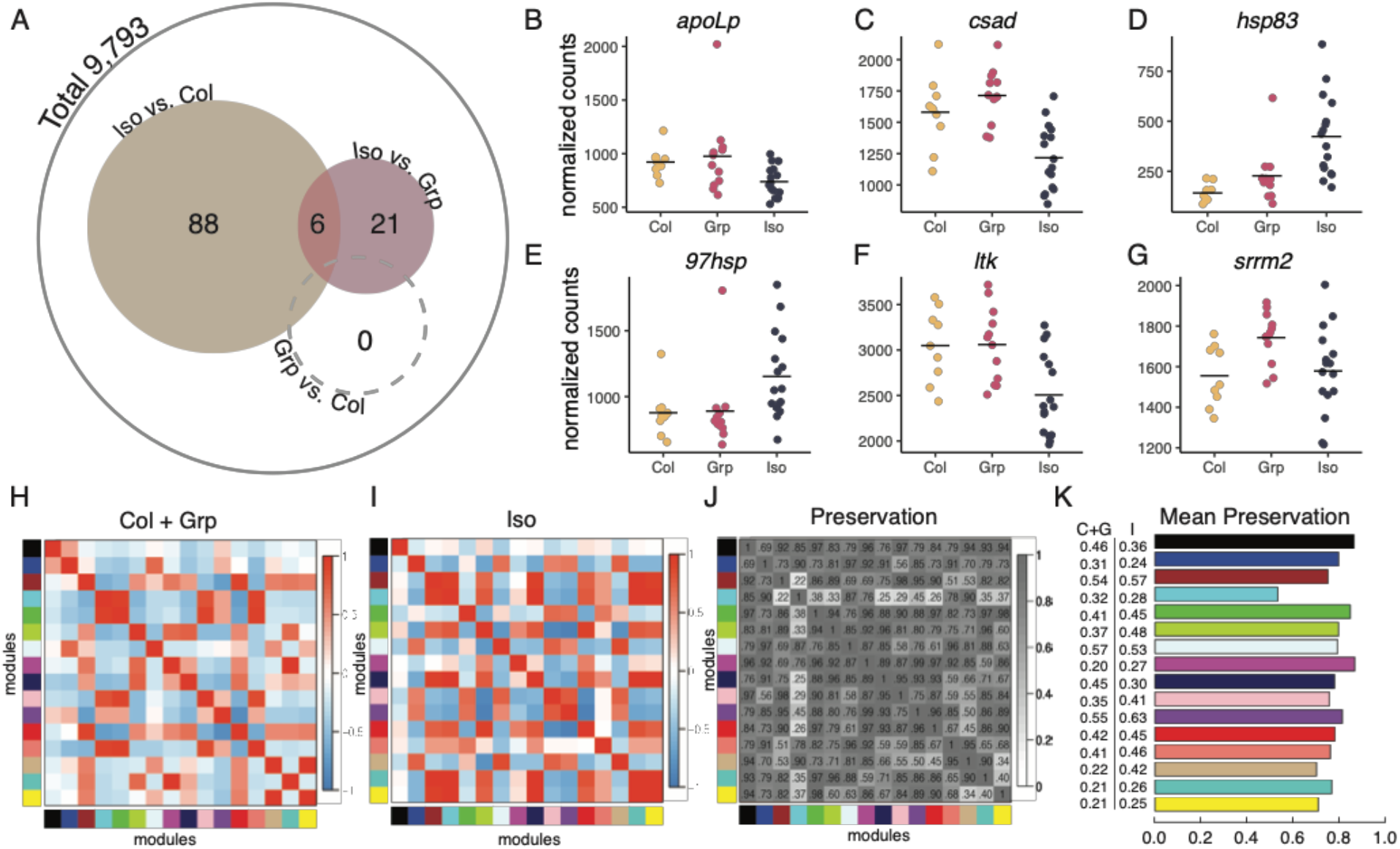
Social isolation disrupts bumblebee neurogenomic landscape. **A.** Differential gene expression analysis. The expression of 94 genes was significantly different between isolated and colony-reared bees. The expression of 27 genes was significantly different between isolated and group-reared bees. Venn diagram shows 6 genes that overlap between these two sets. No genes were differentially expressed between group- and colony-reared bees. **B-G**. Normalized counts of 6 genes in the overlapping region. *apoLp*: apolipophorins, *csad*: cysteine sulfinic acid decarboxylase; *hsp83*: heat shock protein 83; *97hsp*: 97kDa heat shock protein; *ltk*: leukocyte tyrosine kinase receptor; *srrm2*: serine/arginine repetitive matrix protein 2. **H**. Eigengene network headmaps for colony- and group-reared bees. See Methods for details. **I**. Eigengene network heatmaps for isolated bees. See Methods for details. For full module membership, see Table S3. **J**. Heatmap showing preservation between the two networks (1-absolute difference of the two eigengene networks). Darker cells indicate stronger preservation. **K**. Inter- and Intra-module relationships. Barplot showing mean preservation of relationships for each eigengene between colony- and group-reared and isolated bees (inter-module relationships). Numbers indicate mean intra-module correlation within the colony- and group-reared (C+G) and isolated (I) data sets.

In pairwise comparisons, brain transcriptomes of isolated bees showed distinct differences from those of group- and colony-reared bees. We found strong differences in brain gene expression between isolated and colony-reared bees (94 DEGs, FDR < 0.05) and modest differences between isolated and group-housed bees (27 DEGs, FDR < 0.05) (Figure 3A, see Table S1 for full list of genes). Overall, most DEGs showed decreased expression in isolated bees as compared to either of the other two rearing conditions (Table S1). Surprisingly, no DEGs were identified between group and colony-reared bees (Figure 3A). A GOterm enrichment analysis demonstrates that social isolation impacts molecular systems important to social communication, including steroid biosynthesis and signaling processes (Table S2). Overall, our transcriptomic data shows that, much like the changes we observed in behavior, complete social isolation induces significant changes in the expression of key neuromolecular systems important for social living while group-rearing does not significantly alter whole-brain gene expression.

In addition, two genes that are common to the set of isolated vs. group-reared DEGs and the set of isolated vs. colony-reared DEGs (Figure 3B-G) participate in the signaling of juvenile hormone (JH): *apolipophorins* (*apoLp*, Figure 3B) and *heat shock protein 83* (*hsp83*, Figure 3D). In social insects, JH signaling serves as a crucial regulator of reproductive differentiation and social behavior^45–50^. ApoLp and hsp83 are among the suite of proteins that orchestrate the transport of JH to its sites of utilization and initiate its downstream effects^48,51^. In addition, heat shock proteins have previously been identified as key conserved members of the neurogenomic response to social challenge in the honey bee, mouse, and stickleback^52^. The differential expression of both *apoLp*, *hsp83*, and *97hsp* in our transcriptomic data sets strongly suggests that social isolation disrupts signaling of JH in bumblebees, highlighting the importance of this pathway in regulating social behaviors across insects^53^.

Whereas differential gene expression considers each gene individually, network analysis provides insight on the global network properties of the transcriptome. To better understand the gene expression differences across treatment groups, we investigated the gene network dynamics using weighted gene co-expression network analysis (WGCNA^54^). Because there was no evidence for significant differences in gene expression between the colony- and group-reared bees, we combined RNA sequencing data from both groups into a single set for this analysis. We constructed global co-expression networks using data from isolated bees and the combined set of group and colony-reared bee data, then identified modules of genes with linked co-expression (Figure S3, Methods).

To establish concordance and divergence in the network organization between isolated and socially-experienced bees, 16 consensus modules were derived from the weighted average of the two correlation matrices from each behavioral background (Figure 3, Figure S3, Table S3, Methods). We then examined the correlation matrices amongst all gene modules in the isolated and socially-experienced bees (inter-module connectivity, Figure 3H-K). This reveals dysregulation of isolated bee transcriptomes at the module level: genes in the light blue module, for example, show low preservation (negative correlation) in its adjacency relationships to other modules in isolated (I) compared to the colony- and group-reared (C + G) bees (Figure 3J, Table S3). In this module, the mean correlation in colony- and group-reared bees is 0.32, while the mean correlation in isolated bees is 0.28 (Figure 3K). Within all modules, genes in modules from the isolated bee data set tended to show higher connectivity than those in the colony- and group-reared data set (Figure 3K). Interestingly, stronger inter-module connectivity is also a feature of neuronal gene networks in mouse models of autism spectrum disorder^55–57^. Together, our data reveal that social isolation leads to both intra- and inter-module dysregulation of the brain transcriptome (Figure 3H-K).

Given the influence of isolation on brain gene expression, we next interrogated whether social isolation causes broad changes in brain development. The early developmental period in bumblebees is marked by changes in neuropil volume, which reach an adult state around 9 days after eclosion^33^. This maturation process is likely influenced by diverse processes such as learning, endogenous hormone signaling, and experience, including social experiences^58–60^. To determine how the social rearing environment may impact the development of the bumblebee brain, we created an annotated brain template using the full confocal imaging stack of a representative worker bee brain (Figure 4A-B, Video S3). Individual confocal stacks of experimental bees (isolated n=20, group-reared n=24, colony-reared n=22; step size 2.542 um) were fitted to this template for volumetric analysis, and voxels to neuropil regions of interest were summed. To account for individual variation in brain size, we divided the voxels in neuropils of interest by all measured voxels in the brain sample to derive a volume fraction for each bee (Figure 4C-F).

**Figure 4.**
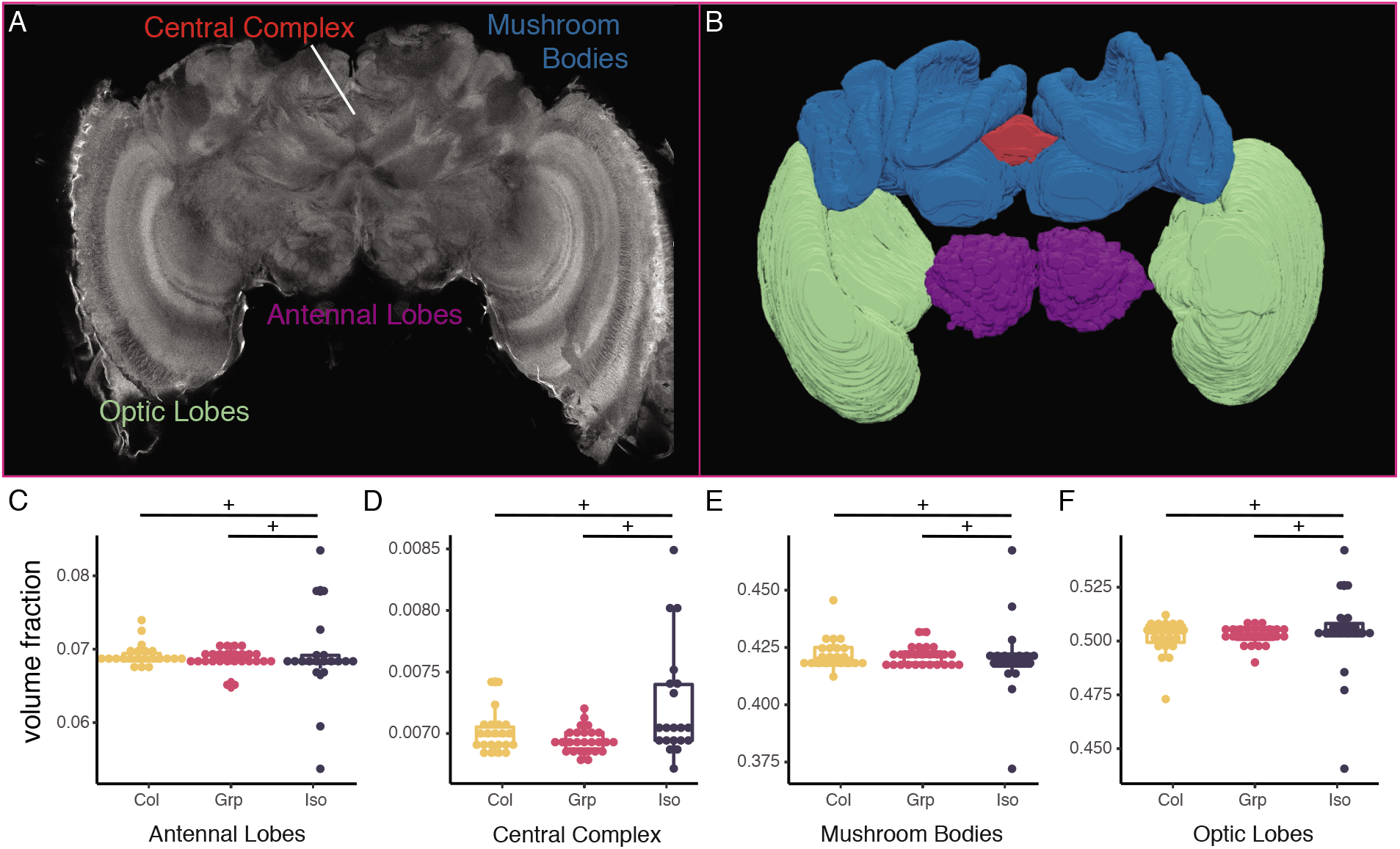
Social isolation destabilizes development of the bumblebee brain. **A**. Confocal slice from the median volume brain used to generate volumetric bumblebee brain atlas. Neuropil label colors correspond to segmented regions in B. For full annotation, see Video S3 and Data Availability. **B**. Segmented volumetric brain atlas. Antennal lobes: purple; Central complex: red; Mushroom bodies: blue; Optic lobes: green. **C-F**. Neuropil volume fractions (raw voxels in area of region/ total voxels). In all regions, the variance was significantly different between the brains of isolated bees and that of group- and colony-reared bees (Fligner-Killeen test, + indicates p value < 0.05).

We find that, while the mean volume fractions of all brain regions were similar across treatment conditions, the variances of the volume fractions were significantly different (Figure 4C-F). For all neuropil regions, inter-individual variance is low amongst colony- and group-reared bees, indicating homogeneity across individuals, but this variance is high in isolated bees (Figure 4C-F). The concordance between the brains of group- and colony-reared bees, and the increased variation in the brains of isolated bees, strongly implies that the social environment acts as a powerful buffering force on the development of the bumblebee. In the complete absence of social cues, the brain may become vulnerable to decanalization, defined as deviations from an optimized phenotype^61–63^. In other words, it appears that social isolation destabilizes and stochastically changes the developmental trajectory of the brain, leading to the greater variation in neuropil volumes observed. This increased variation may be mediated by changes in gene expression or gene network relationships.

Taken together, our results demonstrate that the early life social environment shapes adult behavior in bumblebees, and that these effects are most prominent in social contexts. While all three cohorts had different behavioral repertoires in solo assays, the presence of a social partner led to unique behavioral responses in isolated bees that neither group- nor colony-reared bees exhibited. Isolated bees also demonstrated greater inter-individual variance in behavior (antennae-to-antennae contact) and physiology (neuropil volumes) than bees with any degree of social experience. These differences may be a signature of reduced social competency similar to those described in vertebrates^7–9^. Whether these behavioral changes are due to perturbations in the ability to detect social cues, process the salience of relevant cues, or produce appropriate cues remains unknown. Our study lays the foundation for future research that directly assesses these potential causes as well as the costs of social isolation.

## Acknowledgements

We are thankful to Andrew Webb and Nathaniel Tabris for analytical assistance; Thomas Pisano, Sam Wang, and Gary Laevsky for imaging assistance; Elena Filippova and the Ayroles Lab for reagents; the McBride lab for use of equipment; John Stowers (Loopbio) for technical assistance; and Ian Traniello, Beryl Jones, Sama Ahmed, Dee Ruttenberg, and James Crall for improving the manuscript. J.W.S. acknowledges funding from NIH R01 NS10489 and S.D.K. from NIH 1DP2GM137424-01. In addition, this work was supported by NIH U19 NS104648, the Princeton Catalysis Initiative, and the Lewis-Sigler Institute for Integrative Genomics, and in part by the National Science Foundation, through the Center for the Physics of Biological Function (PHY-1734030).

## Author Contributions

Conceptualization, Z.Y.W., J.W.S., and S.D.K; Methodology, Z.Y.W., G.C.M-S., T.P., J.W.S., and S.D.K.; Investigation, Z.Y.W., G.C.M-S., W.L., H.J.C., T.P., Z.D.; Formal Analysis, Z.Y.W., G.C.M-S.; Resources, J.W.S. and S.D.K; Writing–Original Draft, Z.Y.W.; Writing– Review & Editing, Z.Y.W., G.C.M-S., J.W,S., S.D.K.; Visualization, Z.Y.W.; Supervision, Z.Y.W., G.C.M-S., J.W.S., S.D.K.; Funding Acquisition, J.W.S. and S.D.K.

## Data Availability

All transcriptomic data is deposited in the NCBI SRA database under BioProject ID PRJNA787650. Brain segmentation data is available at https://github.com/kocherlab/BumblebeeIsolation.

## Declaration of Interests

The authors declare no competing interests.

## Materials and Methods

### Animals

Commercial colonies of common eastern bumblebees (*Bombus impatiens*, n=7) were purchased from Koppert Biological Systems (Howell, MI, USA) between June-September 2019. Upon arrival, colonies were visually inspected for the presence of a queen. If no queen was found, or if multiple foundresses were present, colonies were excluded from the study. Colonies were maintained in their original packaging under red light in a room with ambient temperature of 23°C.

### Callow collection

New callows (n = 414) were collected from colonies every morning between 8:30-10:30am. Colonies were chilled at 4°C for 30-45 minutes, at which point the bees were inactive enough to ensure safe removal of callows. Callows were positively identified by their silver-white pigmentation and slow, sluggish gait^28^ (Figure 1A).

### Rearing conditions

Callows were divided into one of 3 rearing conditions, marked accordingly on the dorsal cuticle of the thorax with a paint pen (Sanford Uni-paint, SAN63721), then introduced into their new growth chambers or back into their natal colonies. Total duration of rearing for all bees lasted 9 consecutive days. Isolated bees (Iso) were housed in custom-designed plastic Tritan chambers (7.9 × 5.6 × 3.4 inches) lined with commercial beeswax (UBF10, Betterbee, Greenwich, NY, USA). Each chamber was supplied with HEPA- and carbon-filtered air to eliminate chemical cues and was sound-dampened with anti-vibration padding to remove auditory cues. Each chamber had a feeder (byFormica, B07D5M6F4B) with 40% honey water mixture, which was replaced every other day. Group-reared bees (Grp) were housed in the same chambers with 3 other age-matched nestmates. Each chamber had 2 honey-water feeders, which were replaced every other day. Colony-reared bees (Col) were returned to their natal colonies after marking and left alone for the duration of the 9-day rearing period. Group- and colony-housed individuals were marked on their dorsal thoraces to enable later visual identification. On the 10^th^ day, colony-reared bees were retrieved for subsequent analyses by chilling their natal colonies at 4°C for 30-45 min.

### Behavioral assays

On the 10^th^ day post-eclosion, between 09:00-16:30, experimental bees were removed from their respective rearing chambers and paired with an animal of either the same developmental background (Iso × Iso, n = 24; Grp × Grp, n = 30; Col × Col, n = 28), a different developmental background (Iso × Grp, n = 14; Iso × Col, n = 29; Grp × Col, n = 11; Grp A × Grp B, n = 22), or by themselves (Iso, n = 29; Grp, n = 37; Col, n = 31) in an open Petri dish arena (VWR, 25384-324) lined with commercial beeswax (UBF10, Betterbee, Greenwich, NY, USA). Bees were allowed to interact and move freely within the dish for 30 minutes. Recording started within one minute of introduction to the open arena. Video was captured from above with a Flir Blackfly camera (BFS-U3-32S4M-C) (100 fps, 2048 × 1536 × 1 frame size) on a custom-built Linux computer running LoopBio Motif software. Illumination was provided by 2 infrared panels on the left and right sides of the camera. Each bee was assayed only once. A subset of these bees was collected for subsequent experiments, including RNA sequencing and imaging (see below).

### Body part tracking

SLEAP was used to estimate the pose of the bees and track movement throughout the entire behavioral trial^36^. Twenty-one body parts were labeled to formed a skeleton for pose estimation: head, thorax, abdomen, distal tips of antennae (left antenna 1, right antenna 1), antennal pedicels (left antenna 2, right antenna 2), distal tips of the first wing pair (left wing, right wing), the femur-tibia joint of each leg, and the tarsus of each leg (Figure 1B). We labeled 966 frames with 2 bees from a representative sample of 18 behavioral recordings for a total of 1604 instances. Both bees were always visible, and were often overlapping or partially occluded. Occluded body parts were not labeled. To infer bee tracks across frames, top-down and centroid networks were trained within the sleap.ai framework (TD, ResNet50). Training and inferencing were conducted on a local workstation equipped with an Intel Core i7-5960X CPU, 128 GB DDR4 RAM, NVMe solid state drives and a single NVIDIA Quadro P2000 GPU, or on Princeton University’s High-Performance Computing cluster with nodes equipped with NVIDIA P100 GPUs. Tracks were proofread custom Kalman filter script. Manual adjustments were made to correct any instances in which tracks were swapped between bees.

### Behavior/body posture embedding

To quantify the behavior of the bees, we used the MotionMapper technique^35^. We egocentrized body part traces generated by SLEAP to the thorax body coordinate and thorax-head axis of each bee. In order to have an instantaneous representation of postural dynamics, we performed a continuous wavelet transform on the body part position time series on 25 exponentially spaced frequencies between 0.5Hz and 10Hz, which we empirically determined to be the relevant range for these data. The resulting concatenated spectral densities thus contained information on the power in each of these 25 frequencies for the x and y coordinates of each body part for the length of every trial, so that the postural dynamics of each bee could be described by a 975 element vector for every frame of the video.

In order to define discrete behaviors, we created a low-dimensional representation of these vectors to highlight features of interest. We used t-distributed stochastic neighbor embedding (t-SNE) to embed the concatenated spectral density vectors into a two-dimensional space. t-SNE has the useful property that local similarities will be preserved, such that spectral density vectors that are similar to each other will map onto nearby points in this space, while more global similarities are less important. In order to ensure that we are sampling across all relevant dynamics - that is, that we include even rarely seen dynamics in the spatial embedding - we importance sample across our dataset by first generating a t-SNE embedding of all timepoints for each individual. We then segment this embedding using a watershed transform into 100 different regions, and select points evenly across those regions to contribute to the master t-SNE embedding containing sample points from every trial. Once the master embedding is generated from these samples, we re-embed the non-sample points into the resulting space using the Kullback-Leibler divergence as a distance function. We display the final 2-dimensional ‘space’ as the probability density of the embedding.

We segmented the embedded space of all trials by performing a watershed transform on a less smoothed (sigma=0.5) probability density, with values below a reasonable threshold of probability being ignored, resulting in 38 regions centered around peaks of similar spectral density vectors. We tested several different smoothings and thresholds and chose the one that created a reasonable separation of peaks without over splitting the data. We generated video samples corresponding to each region of various lengths reflecting the varying dwell times of the trajectory of an individual bee’s t-SNE coordinate in each of the regions. By visually inspecting these videos we found these 38 regions, with the exception of region 24, correspond to 5 major stereotyped behavior modalities that we define as: idle (no movement), antennal movement, grooming, locomotion, and a fast locomotion behavior mostly seen in solo trials of group-reared bees (Video S2). Upon visual inspection, video clips from the regions assigned to the fast locomotion state showed bees moving faster than bees than in the locomotion state. Region 24 contained almost the entire t-SNE trajectory of bee #12, and appeared to be the result of idiopathic tracking errors. Bee #12 and region 24 are omitted from the rest of this analysis.

### Compositional analysis of behaviors

#### Paired behaviors

To characterize the effect of isolation on social behavior, we first quantified how frequently paired bees were in close proximity to each other. To do this, we divided the distribution of inter-thorax distances of paired bees by the distribution that would result from random arrangement, allowing us to examine enrichment of specific inter-thorax distances compared to random chance.

#### Defining affiliation

To determine whether distance from a social partner impacts a bee’s overall behavioral repertoire, we quantified changes in the behaviors of paired bees depending on their distance from a social partner. We calculated the Jensen-Shannon divergences (a measure of the difference between probability distributions) between limb dynamics (i.e. the average t-SNE embedded spaces of bees) at different inter-thorax distances in 0.2 cm intervals and the limb dynamics at an inter-thorax distance of 8 cm apart (representative of dynamics at ‘far’ distances)^39^. We accounted for artifacts produced by tracking errors or confined motion by eliminating data from frames in which the bees touched each other or the edge of the arena.

#### Impacts of affiliation

To quantify differences in the behavioral profiles of bees when affiliated versus unaffiliated, we compared the log-ratios of the geometric means of each discrete behavior component to zero. To compare across conditions, we calculated the medians of the distributions of log ratios in behavior components between affiliated and unaffiliated states for each treatment group.

#### Quantification of antennation behaviors

In order to quantify the amount of time that individuals spent contacting other bees with their antennae, we counted the number of antennal touches to different body zones of the social partner. Because different body zones (head, thorax, abdomen, body) have different amounts of available edge space for contact, we normalized the number of antennal touches to body zones of the partner bee by calculating the fraction of the total available edge of the bee each defined zone occupied.

### Tissue collection for RNAseq

Bumblebees were flash frozen on dry ice and stored at −80°C until dissection. To collect the central brain, frozen bees were decapitated with dissecting scissors. With the entire head submerged in RNAlater Ice (Invitrogen, AM7030) over a bed of dry ice/ethanol, large sections of the dorsal and ventral head cuticle and mandibles were chipped away to expose neural tissues. The chipped heads were transferred to a 10x volume of RNAlater-ICE (Invitrogen AM7030) solution, submerged, and allowed to incubate at 4°C for 16 hours before subdissection of the central brain over wet ice. Fat bodies, compound eyes, and ocelli were removed from the head mass. Brains were placed into individual 1.5 ml Eppendorf tubes with a 2.8mm bead (OPS Diagnostics, GBSS 089-5000-11) and homogenized for 10 minutes at maximum speed on a Qiagen TissueLyserII.

### RNA extraction and TM3’-seq

We extracted RNA from whole brain homogenates using the Dynabeads mRNA DIRECT kit (Invitrogen, 61011) according to manufacturer’s protocol with homemade low-salt buffer (LSB, 20mM Tris-HCl[pH 7.5], 150 mM NaCl, 1mM EDTA) and Lysis/Binding Buffer (LBB, 100mM Tris-HCl[pH 7.5], 500 mM LiCL, 10mM EDTA[pH 8], 1% LiDS, 5mM DTT). RNA quality and concentration were checked on a Tapestation in the Princeton Genomics Core before proceeding with library preparation. Samples with RNA integrity scores below 8.8 were excluded from study. We prepared TM3’-seq libraries according to published protocols^44^ using i5 and i7 primers and Tn5 generously provided by the Ayroles lab. Libraries of 43 bees were sequenced on an Illumina NovaSeq instrument (single-end, S1 100nt lane), generating ~450 million reads. Samples were demultiplexed by the Princeton Genomics Core and samples with low read counts (<1 million reads, n = 3) were excluded from study. A total of 16 isolated, 15 group-reared, and 9 colony-reared bees were included for transcriptomic analyses.

### Transcriptomic Data Preprocessing

We followed the recommended pipeline for TM3’seq data processing^44^ (see also https://github.com/Lufpa/TM3Seq-Pipeline). Reads were trimmed with custom trimmers using Trimmomatic^64^. Reads were aligned to the reference *Bombus impatiens* genome (BIMP_2.2, GCF_000188095.3) using STAR^65^. Small reads and duplicated reads were filtered out with SAMtools^66^. Mapped reads were counted using featureCounts^67^. Aggregate data preprocessing results can be viewed in MultiQC v1.8 here: file:///Users/zyanwang/Dropbox%20(Princeton)/RNAseq/multiqc_report_2.html

### Differential gene expression analysis

Analysis of differential genes was performed using the DESeq2 package^68^ in RStudio version 1.2.5001 running R version 3.6.1. The standard DESeq2 workflow was applied to the unique (deduplicated) aligned raw reads, and an additive model was built with source colony and treatment as factors. For pairwise testing of differential expression, colony-reared bees were set as the control and the false discovery rate (FDR) was set at 0.05.

### GOTerm Enrichment

Gene Ontology terms were assigned to genes using Trinotate^69^. Gene set enrichment analysis was performed using the topGO package in R^70^. Only Biological Process terms were considered, and a Fisher’s exact test was used to perform the enrichment test using the “elim” algorithm in topGO. FDR was set at 0.05.

### Weighted gene correlation network analysis

Co-expression analysis of brain RNA sequencing data from all bees was implemented with the WGCNA package in R^54^. Consensus correlation matrices were constructed and converted to adjacency matrices that retained information about the sign of the correlation^71^. Adjacency matrices were raised to a soft power threshold of 10. This was empirically determined based on a measure of R2 scale-free topology model fit that maximized and plateaued over 0.8. The soft-power thresholded adjacency matrices were converted into a topological overlap matrix (TOM) and a topological dissimilarity matrix (1-TOM). We then performed agglomerative hierarchical clustering using the average linking method on the TOM dissimilarity matrix. Gene modules were defined from the resulting clustering tree, and branches were cut using the hybrid dynamic tree cutting function: the module detection sensitivity (deepSplit) was set to 2 (default), minimum module size 30 (default), and the cut height for module merging set to 0.25 (modules whose eigengenes were correlated above 0.75 were merged). This yielded 16 consensus modules (Figure 3, Figure S3), each assigned a color label. For each gene module, a summary measure (module eigengene) was computed as the first principal component of the module expression profiles. Genes that could not be clustered into any module were assigned to module M0 and not used for any downstream analysis. Correlation matrices for module eigengenes were then calculated separately for each data set (i.e. we considered RNA sequencing data from colony- and group-reared bees as the first data set, and isolated bees as the second data set) for comparison.

We also constructed set-specific modules in order to relate network relationships unique to each data set to the global relationships in the consensus modules. Network construction and module detection was performed as described above. We related set-specific modules to consensus modules by calculating the overlap of each pair of modules and using Fisher’s exact test to assign a p-value to each of the pairwise overlaps (Figure S3).

### Whole mount dissections and tissue preparation for confocal imaging

To measure neuropil volumes of bumblebees, we created a brain atlas based on confocal image stacks of the bee brain’s natural autofluorescence. Bees were anesthetized with CO2 and decerebrated. Mandibles were removed, then heads were placed in fresh 4% paraformaldehyde (PFA) at 4°C overnight, rocking. A top-down photograph of the bee head was taken, and head width was measured in Fiji. The brains were subdissected in cold PBS and fixed in PFA at 4°C overnight. The next day, brains were washed in fresh phosphate-buffered saline (PBS) 3 × 10 min, transferred to a glass scintillation vial, and post-fixed in 2% glutaraldehyde at 4°C for 48 hrs. After post-fixation, brains were washed 3 × 10 min in PBS, submerged in formamide bleach (76% PBS, 20% 30% H2O2, 1% 10% Triton-X100, 3% formamide) for 75 min at room temperature, and washed again 3 × 10 min in PBS. Brains were then dehydrated in ethanol: 1 × 10 min washes of 30%, 50%, 70%, 90%, and 95% EtOH, then 3 × 10 min washes of 100% EtOH. Samples were stored in 100% EtOH until clearing and imaging. Brains were cleared in methyl salicylate (Sigma Aldrich, M2047) for 30 min at room temperature, then mounted in fresh methyl salicylate on a glass slide for confocal imaging.

### Confocal imaging and brain atlas construction

All imaging was performed in the Princeton Confocal Imaging Facility on a Nikon A1 laser confocal microscope and a PC machine running the Nikon Elements Software package. Samples were scanned in the 488 nm laser line. Images were optically sectioned at 2.542um until the entire brain was imaged in series at 10x magnification. Large image grab was used to image the entire field of view in 4 quadrants, then stitch quadrants together to create a single 1895 × 1895 image. The following regions of the reference worker brain was manually segmented based on visible boundaries visualized with autofluorescence using a Wacom drawing tablet and the segmentation/3D reconstruction software ITK-SNAP: the central complex (including protocerebral bridge and nodules), antennal lobes, mushroom body and mushroom body lobes, and optic lobes were manually segmented. This reference brain was used as the template for downstream brain registration.

### Measuring brain volumes

The elastix package^72^ was used to register confocal images of experimental brains to the template brain. The Jacobian determinants were calculated using transformix, and after transformation, voxels corresponding to each neuropil region were summed. Voxel data was plotted in RStudio using the ggplot2 package. The Fligner-Killeen test of homogeneity of variances was used across samples (p-value < 0.05).

## Supplementary Figure Legends

**Figure S1. Extended behavior analysis**

**A.** Watershed transform of the embedded space of body dynamics showing the 38 regions identified around separate density peaks. By visually inspecting video clips from each region, we grouped regions together based on similar stereotyped behaviors, indicated by the bold black lines. Region 24 was excluded since it contained the dynamics of a single bee. **B-D**. Probability density maps showing the distribution of timepoints from the solo trials of isolated (**B**), group-reared (**C**), and colony-reared (**D**) bees. **E-G**. Histograms showing the different thorax speed distributions of solo assayed isolated, group-, and colony-reared bees. **H**. Behavior compositions for all trial types. For mixed pairings, the treatment condition noted first is the one displayed (e.g. isoxgrp indicates data from isolated bees that have been paired with group-reared bees).

**Figure S2. Extended affiliation analysis**

**A-G**. Inter-thorax occupancy shown as enrichment over the null model of random chance for all paired trials, as labeled. Data from the homogeneous pairings are also presented in the body of the paper, and are shown here for completeness. **H-I**. Behavior compositions for affiliated (**H**) and unaffiliated (**I**) bees for all paired trials. For mixed pairings, the treatment condition noted first is the one displayed (e.g. isoxgrp indicates data from isolated bees that have been paired with group-reared bees).

**Figure S3. Weighted gene correlation network analysis.**

A. Consensus gene dendrogram obtained by clustering the dissimilarity of genes from all samples (Colony-, Group-, and Isolation-reared) based on consensus topological overlap (see Methods). Corresponding module colors plotted below. B. Dendrogram of consensus module eigengenes in colony- and group-reared bees. C. Dendrogram of consensus module eigengenes in isolated bees. D. Correspondence of modules built from colony- and group-reared bee data (y-axis) and consensus modules (x-axis). Numbers in the table indicate gene counts in the corresponding module. Cell color indicates −log(p), where p = Fisher’s exact test p-value for the overlap of the two modules: the more significant the overlap, the redder the cell. E. Correspondence of modules built from isolated bee data (y-axis) and consensus modules (x-axis).

**Figure S4. Brain voxel measurements.**

**A**. Head width of bumblebees. Variances are not significantly different from each other (Levene’s test); means are not significantly different from each other (Kruskal-Wallis test). **B**. Total raw voxels. Variances are not significantly different from each other (Levene’s test); means are not significantly different from each other (Kruskal-Wallis test). **C-E**. Normalized volumes of brain regions by treatment. Samples plotted by increasing total volume along the x-axis. CC: central complex, AL: antennal lobe, MB: mushroom bodies; OL: optic lobes

## Supplementary Items

Video S1. SLEAP-tracked pair of bees

Video S2. Discrete behavior map examples

Video S3. Worker Bee Brain Reference Template

Table S1. Table of Differentially Expressed Genes

Table S2. Table of GOTerm Enrichment lists

Table S3. Table of WGCNA Module Membership

## Notes

https://github.com/kocherlab/BumblebeeIsolation

### Competing Interest Statement

The authors have declared no competing interest.

