## Supplemental Figures for "Isolation disrupts social interactions and destabilizes brain development in bumblebees"

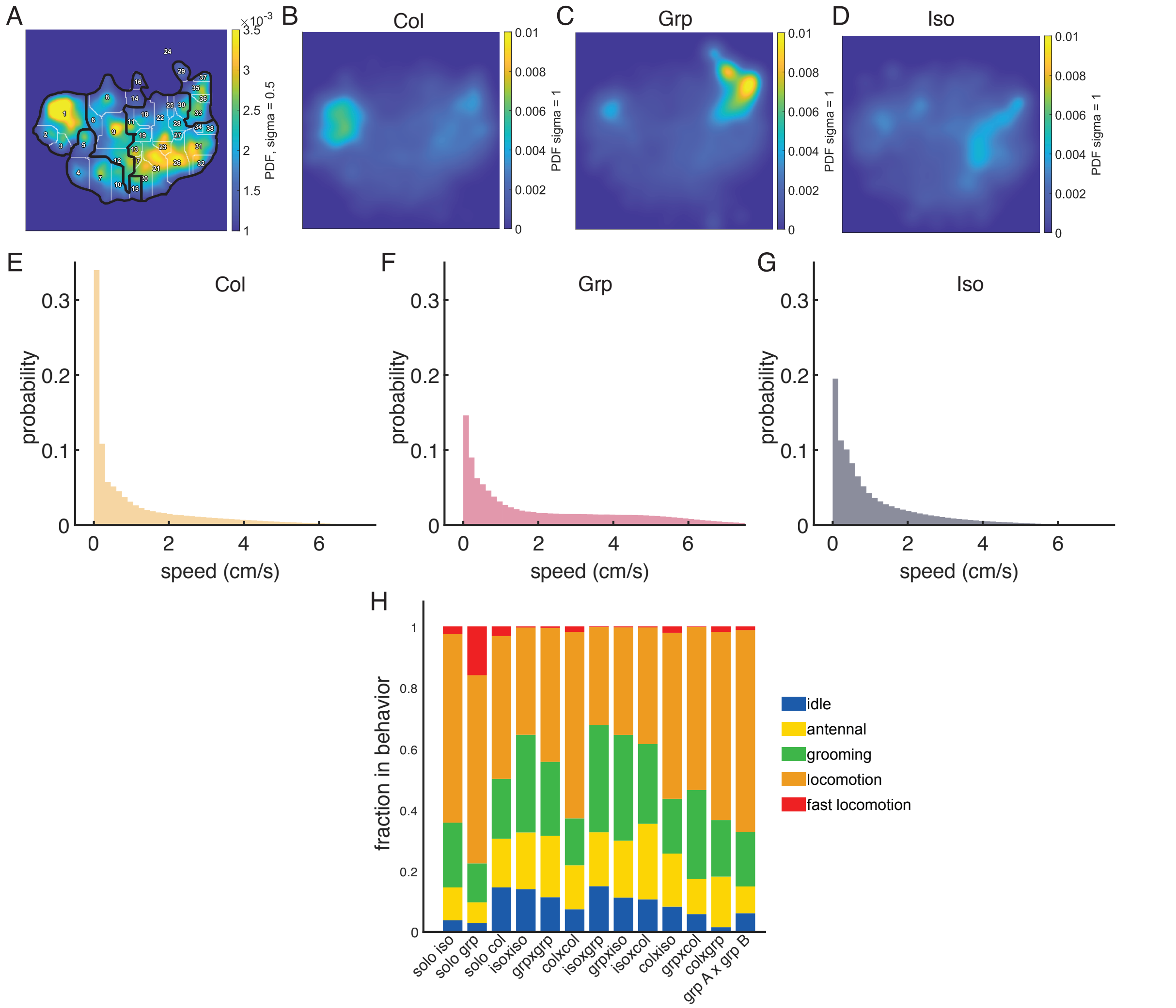


**Figure S1. Extended behavior analysis**

**A.** Watershed transform of the embedded space of body dynamics showing the 38 regions identified around separate density peaks. By visually inspecting video clips from each region, we grouped regions together based on similar stereotyped behaviors, indicated by the bold black lines. Region 24 was excluded since it contained the dynamics of a single bee. **B-D.** Probability density maps showing the distribution of timepoints from the solo trials of isolated (**B**), group-reared (**C**), and colony-reared (**D**) bees. **E-G.** Histograms showing the different thorax speed distributions of solo assayed isolated, group-, and colony-reared bees. **H**. Behavior compositions for all trial types. For mixed pairings, the treatment condition noted first is the one displayed (e.g. isoxgrp indicates data from isolated bees that have been paired with group-reared bees).

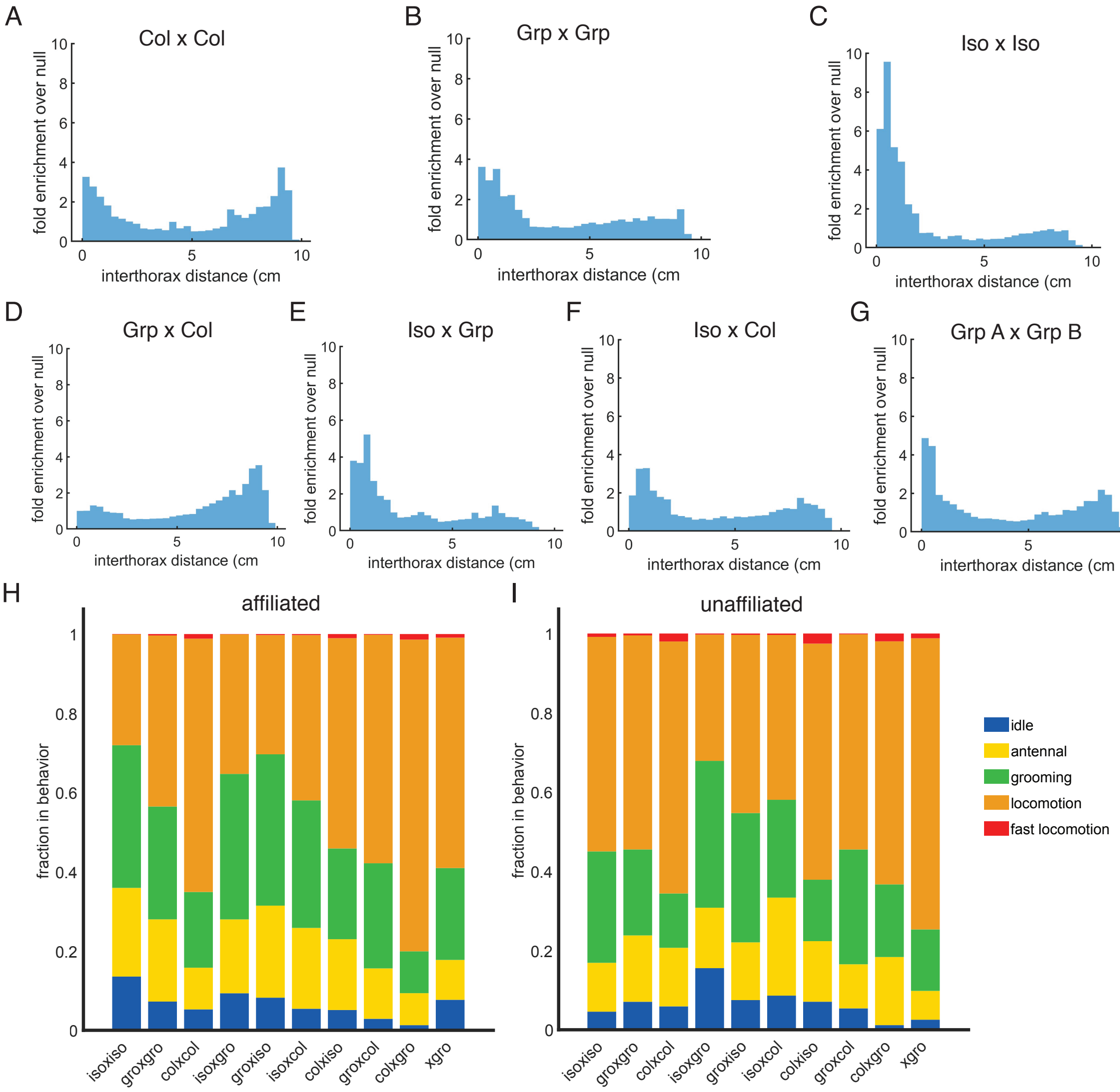


**Figure S2. Extended affiliation analysis**

**A-G**. Interthorax occupancy shown as enrichment over the null model of random chance for all paired trials, as labeled. Data from the homogeneous pairings are also presented in the body of the paper, and are shown here for completeness. **H-I**. Behavior compositions for affiliated (**H**) and unaffiliated (**I**) bees for all paired trials. For mixed pairings, the treatment condition noted first is the one displayed (e.g. isoxgrp indicates data from isolated bees that have been paired with group-reared bees).

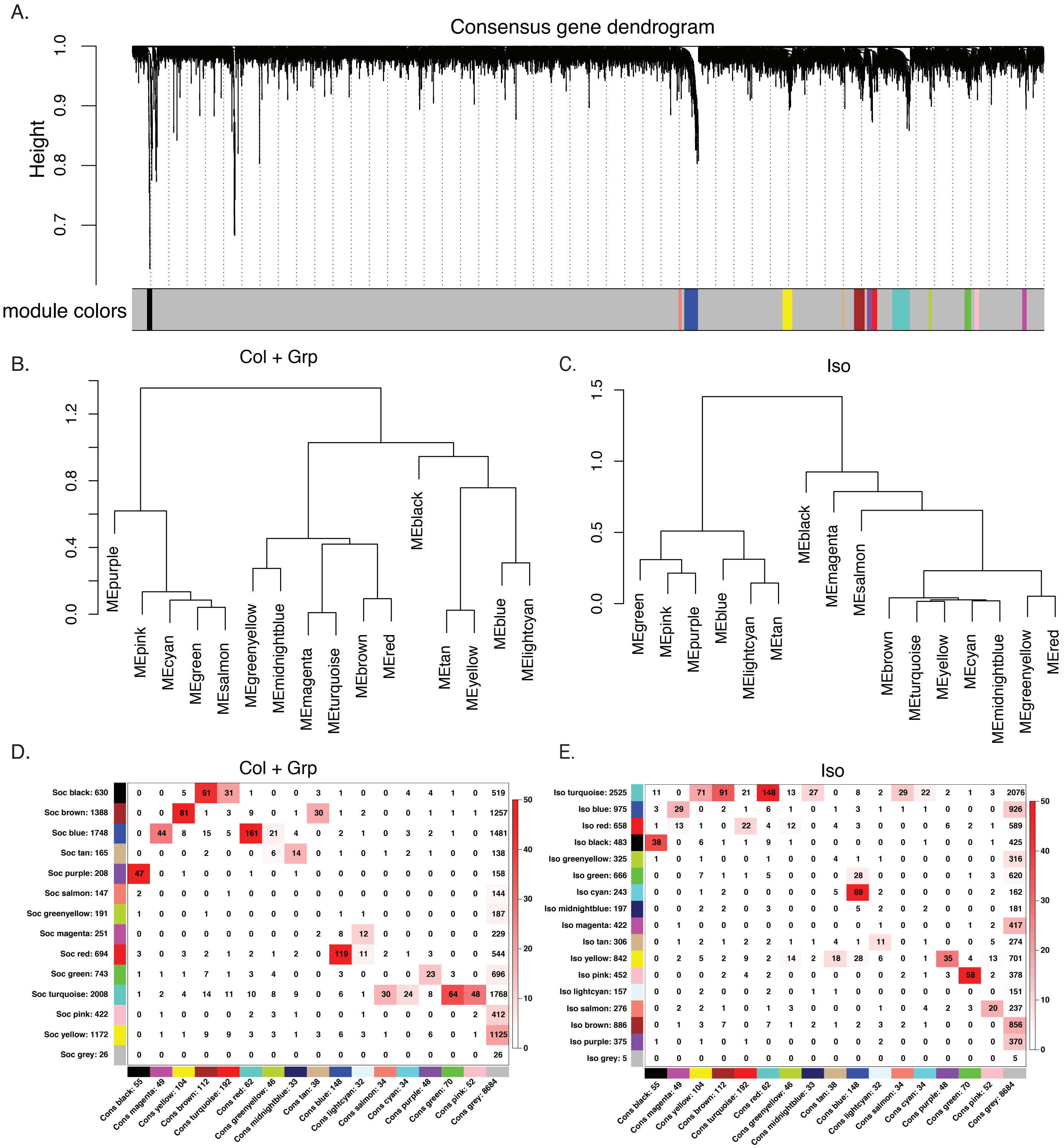


**Figure S3. Weighted gene correlation network analysis.**
**A.** Consensus gene dendrogram obtained by clustering the dissimilarity of genes from all samples (Colony-, Group-, and Isolation-reared) based on consensus topological overlap (see Methods). Corresponding module colors plotted below. **B**. Dendrogram of consensus module eigengenes in colony- and group-reared bees. **C**. Dendrogram of consensus module eigengenes in isolated bees. **D**. Correspondence of modules built from colony- and group-reared bee data (y-axis) and consensus modules (x-axis). Numbers in the table indicate gene counts in the corresponding module. Cell color indicates -log(p), where p = Fisher's exact test p-value for the overlap of the two modules: the more significant the overlap, the redder the cell. **E**. Correspondence of modules built from isolated bee data (y-axis) and consensus modules (x-axis).

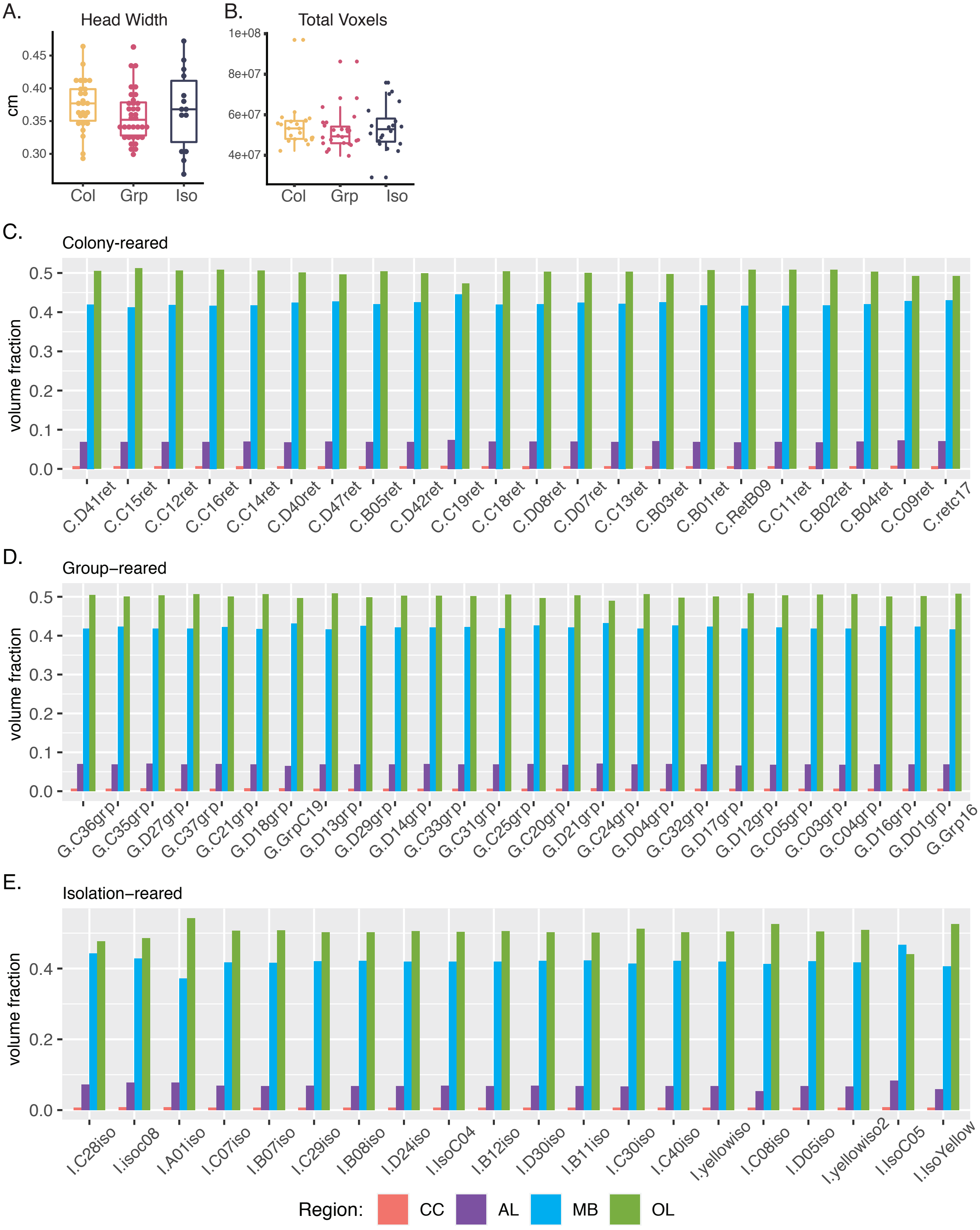


**Figure S4. Brain voxel measurements.**
**A.** Head width of bumblebees. Variances are not significantly different from each other (Levene's test); means are not significantly different from each other (Kruskal-Wallis test). **B**. Total raw voxels. Variances are not significantly different from each other (Levene's test); means are not significantly different from each other (Kruskal-Wallis test). **C-E**. Normalized volumes of brain regions by treatment. Samples plotted by increasing total volume along the x-axis. CC: central complex, AL: antennal lobe, MB: mushroom bodies; OL: optic lobes
